# Application of a dense genetic map for assessment of genomic responses to selection and inbreeding in *Heliothis virescens*

**DOI:** 10.1101/025981

**Authors:** Megan L. Fritz, Sandra Paa, Jennifer Baltzegar, Fred Gould

## Abstract

Adaptation of pest species to laboratory conditions and selection for resistance to toxins in the laboratory are expected to cause inbreeding and genetic bottlenecks that reduce genetic variation. *Heliothis virescens*, a major cotton pest, has been colonized in the laboratory many times, and a few laboratory colonies have been selected for Bt resistance. We developed 350 bp Double-Digest Restriction-site Associated DNA-sequencing (ddRAD-seq) molecular markers to examine and compare changes in genetic variation associated with laboratory adaptation, artificial selection, and inbreeding in this non-model insect species. We found that allelic and nucleotide diversity declined dramatically in laboratory-reared *H. virescens* as compared with field-collected populations. The declines were primarily due to the loss of low frequency alleles present in field-collected *H. virescens*. A further, albeit modest decline in genetic diversity was observed in a Bt-selected population. The greatest decline was seen in *H. virescens* that were sib-mated for 10 generations, where more than 80% of loci were fixed for a single allele. To determine which regions of the genome were resistant to fixation in our sib-mated line, we generated a dense intraspecific linkage map containing 3 PCR-based, and 659 ddRAD-seq markers. Markers that retained polymorphism were observed in small clusters spread over multiple linkage groups, but this clustering was not statistically significant. Here, we confirmed and extended the general expectations for reduced genetic diversity in laboratory colonies, provided tools for further genomic analyses, and produced highly homozygous genomic DNA for future whole genome sequencing of *H. virescens*.

## Introduction

Laboratory-reared insect colonies are important resources for many types of entomological experiments. They are used to quantify physiological or behavioral differences between insect populations or species (Dekker *et al.*, 2006; Dobzhansky & Spassky, 1954; Fritz *et al.*, 2015; Groot *et al.*, 2005; Shaw *et al.*, 2000; Sokolowski, 1980; Tomaru *et al.*, 2000), identify the genetic architecture of insect traits (Gahan *et al.*, 2010, Mackay *et al.*, 2012, Oppenheim *et al.*, 2012), develop insect populations that express desirable traits (Collins, 1984; Goldman *et al.*, 1986; Gould *et al.*, 1995; Hoy, 1990; Pradeep *et al.*, 2005), and generate genetically modified species as a means of pest control (de Valdez *et al.*, 2011). A major concern for researchers maintaining insect colonies is the degree to which adaptation to the laboratory environment affects insect genotypic, and thereby phenotypic diversity (Boller, 1972; Huettel, 1976).

The phenotypic consequences of adaptation to the laboratory depend upon the trait of interest, and range from undetectable to severe (Baeshen *et al.*, 2014; Fox *et al.*, 2007; Gerloff *et al.*, 2003; Raulston, 1975; Roush, 1986). Observed phenotypic changes can be attributed to inadvertent selection for traits that are favorable in the laboratory environment (Roush, 1986), inbreeding depression *(i.e*. reduction in fitness caused by matings between related individuals; reviewed in Charlesworth & Willis, 2009; Mackauer, 1976), or the interaction of the two. Indeed, the selection that occurs during colony establishment creates conditions conducive to inbreeding (Roush, 1986). Families with higher fitness under laboratory conditions contribute disproportionately to the reproductive pool, thereby increasing the probability of matings between related individuals. Where selection is very strong, as in the production of an insecticide resistant colony, measures must often be taken to minimize the effects of inbreeding and thereby inbreeding depression (Gould *et al.*, 1995). Overall, the expectation is that the selection and inbreeding that takes place during insect colonization results in an overall loss of genetic diversity (Munstermann, 1994), and concomitant genome-wide increase in homozygosity (reviewed in Etzel & Legner, 1999).

Previous studies that have examined genetic differences between field-collected, laboratory-adapted, and inbred populations of non-model insects have primarily focused on Dipteran species and were limited to small numbers of molecular markers (Mukhopadhyay *et al.*, 1997; Munstermann, 1994; Norris *et al.*, 2001). Such small numbers of markers allow for estimation of the genome-wide average change in genetic variability across populations, but cannot be used to examine fine-scale patterns of genomic change. Examination of these patterns allows for identification of where and how genetic variation, the raw material necessary for environmental adaptation, is maintained (Dobzhansky & Spassky, 1954). The relatively recent development of high-throughput sequencing combined with reduced-representation DNA library preparation techniques allows for the discovery of hundreds to thousands of new molecular markers, even in species for which genomic data are absent (Davey *et al.*, 2012). Here we used Double-Digest Restriction-Site Associated DNA Sequencing (ddRAD-seq; Peterson *et al.*, 2012), one type of reduced representation library preparation, for *de novo* construction of molecular markers in the non-model species, *Heliothis virescens*.

The tobacco budworm, *H. virescens*, is an historically important pest of cotton throughout much of the Southeastern United States (Blanco, 2012). This non-model Lepidopteran species has been colonized a number of times for investigations of mating and host-selection behaviors (Sheck & Gould, 1995; Sheck & Gould, 1996; Sheck *et al.*, 2006), as well as detecting the underlying genetic basis for insecticide resistance (Gahan *et al.*, 2001; Gahan *et al.*, 2010; Taylor *et al.*, 1993). We used our newly developed ddRAD-seq markers to examine and compare the effects of colonization, selection, and sib-mating on *H. virescens* genome-wide measures of genetic diversity. To examine fine-scale patterns of change in genetic diversity, we also used our ddRAD-seq markers to generate a dense intraspecific genetic map for *H. virescens*. This map consists of 659 high quality 350-bp markers which will serve as an important genomic resource to the entomological community.

Overall, our research aims to:

1. Quantify overall patterns of change in genomic diversity across field-collected, laboratory-reared (non-selected), Bt-selected, and sib-mated *H. virescens*.
2. Determine whether the observed degree of inbreeding in our sib-mated *H. virescens* calculated from our ddRAD-seq genotypic data matched theoretical expectations (Falconer & Mackay 1996).
3. Use ddRAD-enabled linkage mapping to determine if specific genomic regions were resistant to fixation, even under intense inbreeding, by comparing genotypic data from long-term colony and sib-mated lines.

## Results

We sequenced 204 *H. virescens* individuals from a total of 6 populations that were used in a population-level analysis of genomic change associated with laboratory colonization, artificial selection and inbreeding. These populations were comprised of 2 field-collected (LA, TX populations collected in 2012), 2 laboratory-reared (BENZ, YDK), and 1 Bt-selected population (YHD2), as well as specimens from a single inbred family following 10 generations of full-sibling mating (see Table 1 for information on population history, sample sizes, and read counts). Three of these populations, YDK, YHD2, and the inbred line were founded from a collection in Yadkin County, NC, in 1988 (Gould *et al.*, 1995), but were thereafter subjected to different rearing conditions, allowing us to make comparisons of population-genomic change within the same genetic background. In addition, 99 individuals (1 BENZ parent, 1 BENZ-YHD2 hybrid parent and 97 progeny) were sequenced for linkage analysis. This produced a total of 105,487,499 Illumina MiSeq reads (38,221,995 and 67,265,504 for linkage- and population-level analyses, respectively) that passed quality filters (data available upon request).

**Table 1.**
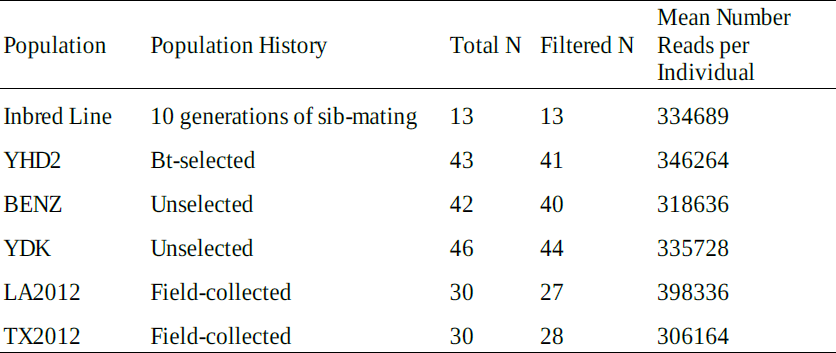
Population history, sample sizes (N) and ddRAD-seq read summary data. Filtered N refers to the population sample size following removal of individuals with low read counts.

| Population | Population History | Total N | Filtered N | Mean Number<br>Reads per<br>Individual |
| --- | --- | --- | --- | --- |
| Inbred Line | 10 generations of sib-mating | 13 | 13 | 334689 |
| YHD2 | Bt-selected | 43 | 41 | 346264 |
| BENZ | Unselected | 42 | 40 | 318636 |
| YDK | Unselected | 46 | 44 | 335728 |
| LA2012 | Field-collected | 30 | 27 | 398336 |
| TX2012 | Field-collected | 30 | 28 | 306164 |

### Genomic diversity among *H. virescens* populations

On average, 338,892 sequencing reads (*s.d*. = 113,397) were produced per individual, and the variation in read count was spread uniformily across populations (Supplementary Figure 1). Ninety-four percent of individuals had read counts between 90,000 and 688,000, and these were fed into the Stacks pipeline (Catchen *et al.*, 2011; 2013) for *de novo* locus construction (Supplementary Figure 1). Loci constructed by Stacks had an average of 6× depth of coverage per individual. In total, 4,281 polymorphic 350-bp ddRAD-seq markers (hereafter loci) were detected in at least one individual per population across all populations. Two well-documented challenges commonly encountered when working with moderate coverage reduced representation library data like ours are: 1) uneven distribution of missing data across sets of loci (Davey *et al.*, 2012; Xu *et al.*, 2014), and 2) under-sampling of heterozygotes (Li *et al.*, 2009; Nielsen *et al.*, 2011). Both reduce confidence in final genotypes called by genotyping-by-sequencing SNP calling algorithms, including the algorithm used in Stacks. To overcome these challenges, we examined several subsets of these 4,281 polymorphic loci for our downstream population genomic analyses. These subsets contained between 125-1231 loci, and were chosen based upon the overall proportion missing genotype calls in the subset. The smallest subset consisted of loci for which over 75% of individuals per population had genotypic data present and were therefore likely sequenced to greatest depth of coverage. Each larger subset allowed additional loci at the expense of coverage *(i.e*. more missing genotypic data were allowed; Supplementary Table 1). By using multiple datasets, we were able to examine whether the presence of missing genotype calls influenced overall estimates of genomic diversity across populations.

We first examined all subsets of loci and determined the mean and maximum numbers of alleles per locus (Supplementary Table 1). In our case, alleles were not analogous to SNPs, but rather the accumulation of SNPs per 350 bp locus per individual. For the total sequenced population (n = 192 total *H*. *virescens*), the mean numbers of unique alleles detected per locus ranged from 29 to 34 depending upon the number of loci included in the analysis. As more loci were included, the average number of unique alleles detected per locus decreased. However, the maximum number of unique alleles detected in the total population increased from 86 in the smallest subset of loci to 94 in the 3 larger subsets. We also examined the proportion of loci that were fixed (*i.e*. only a single allele present) across populations. Across subsets, few loci were fixed for a single allele in laboratory-reared (5.6-10.9%), Bt-selected (5.3-7.3%) and field-collected (0-2.4%) populations (Supplementary Table 1). Yet over 80% of loci were fixed in the inbred line following 10 generations of sib-mating (Supplementary Table 1). Of the 125 loci with the fewest missing genotype calls, 86% were fixed in the inbred line. Expanding the number of loci to include those with more missing genotypes (n= 378, 573, 1231) reduced the percentage of fixed loci in the inbred line by up to 5% (Supplementary Table 1).

We then determined the mean number of unique alleles present per locus for each subset of loci (Supplementary Figure 2). In general, we found no within population differences in the mean numbers of unique alleles detected among subsets of loci, and therefore we used a single, conservatively chosen subset of loci (n = 378) where at least 10 individuals were genotyped per population per locus for further analysis. The mean numbers of unique alleles per locus were 2.1 for the inbred line, 5.3 for the Bt-selected population, 5.4 and 4.4 for the non-selected, laboratory-reared populations (YDK and BENZ, respectively), and 18.4 and 17 for the field-collected populations (LA and TX, respectively). However, our sample sizes (*i.e*. numbers of individuals sequenced; see Table 1) differed for each population, and it was unclear whether differences between the aforementioned means were caused by sample size or population-level differences. For example, it is to be expected that as sample size increases there will be an increase in the probability of sampling additional, likely rare, alleles. Therefore, we randomly sub-sampled pools of alleles 6, 12, 18, and 24 times without replacement for each population. This allowed us to hold sample sizes constant across populations, and infer whether the mean numbers of unique alleles truly differed by population. As expected, we found that increasing the total number of alleles sampled led to an increase in the mean numbers of unique alleles per locus for all but the inbred line (Figure 1). Yet we also found strong population-level differences. Regardless of the number of alleles sampled, field populations always exhibited the greatest allelic diversity, followed by selected and non-selected colony populations. The lowest allelic diversity was observed in the inbred line.

**Figure 1.**
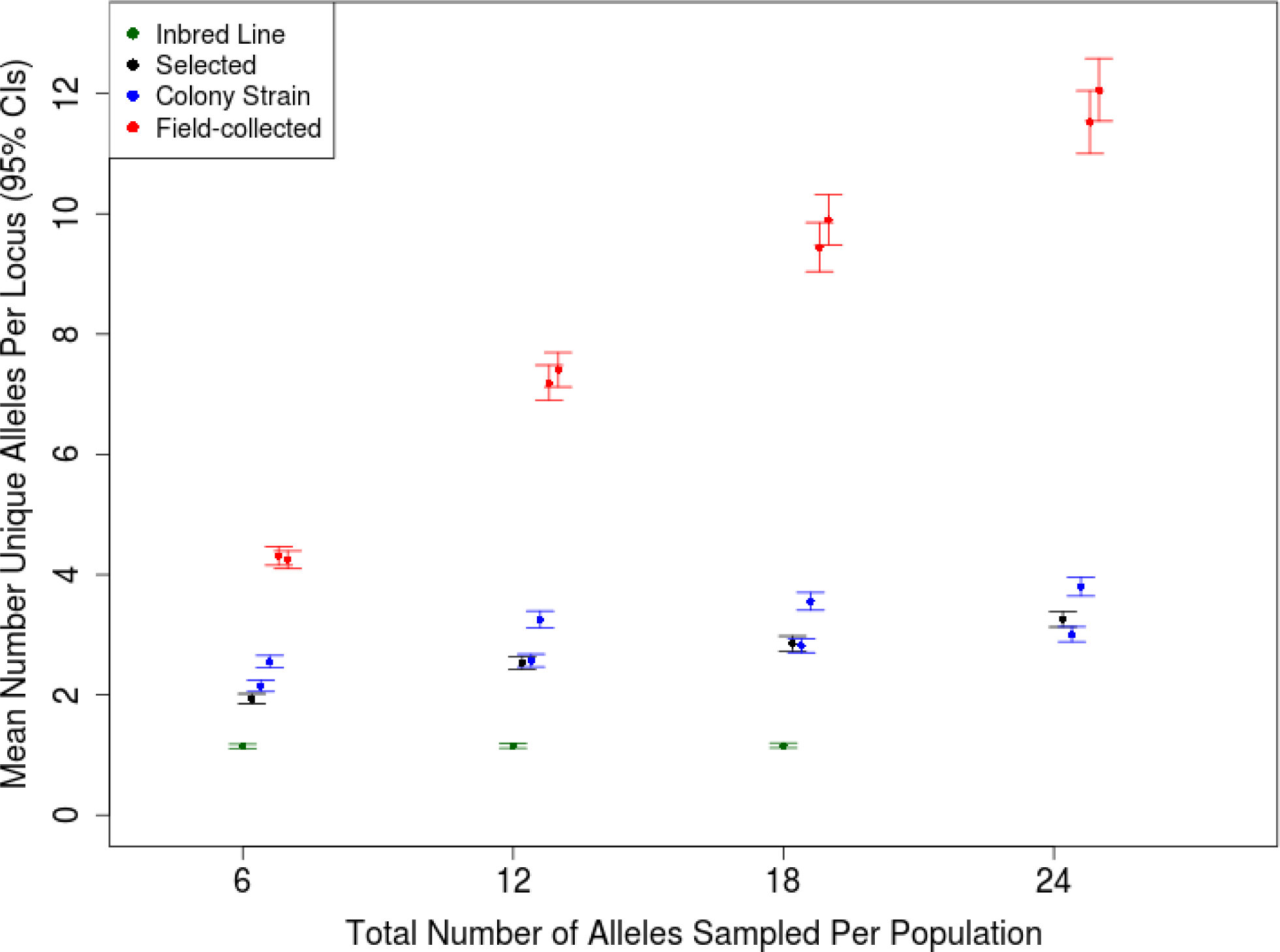
Mean numbers of unique alleles detected among 378 loci depends upon the number of alleles sampled per population. For all but the inbred line (in green), as sample size increases, so does the number of unique alleles detected per locus. Due to low survivorship in the inbred line, no mean was computed for an allelic sample size per locus of 24.

When 18 alleles were randomly sampled per population per locus, we detected an average of just over 1 unique allele per locus in the inbred line, indicating that most loci were fixed for a single allele. For the inbred line, 52 of the 378 loci did not reach fixation. Of these, forty-seven had 2 unique alleles, four had 3 unique alleles, and one had 4 unique alleles when 18 were randomly sampled. On average, Bt-selected and non-selected colony strains each had *ca*. 3 unique alleles per locus, and field-collected populations had *ca*. 9 unique alleles per locus (Figure 1). The majority of unique alleles present in the field-collected populations (70.3% and 68.7% for LA and TX populations, respectively) were observed only once (of 18 alleles; Figure 2). Such low frequency alleles were less common in the Bt-selected and non-selected laboratory populations (35.1% and 21.5%, respectively) and rare (0.7%) in the sib-mated inbred line (Figure 2).

**Figure 2.**
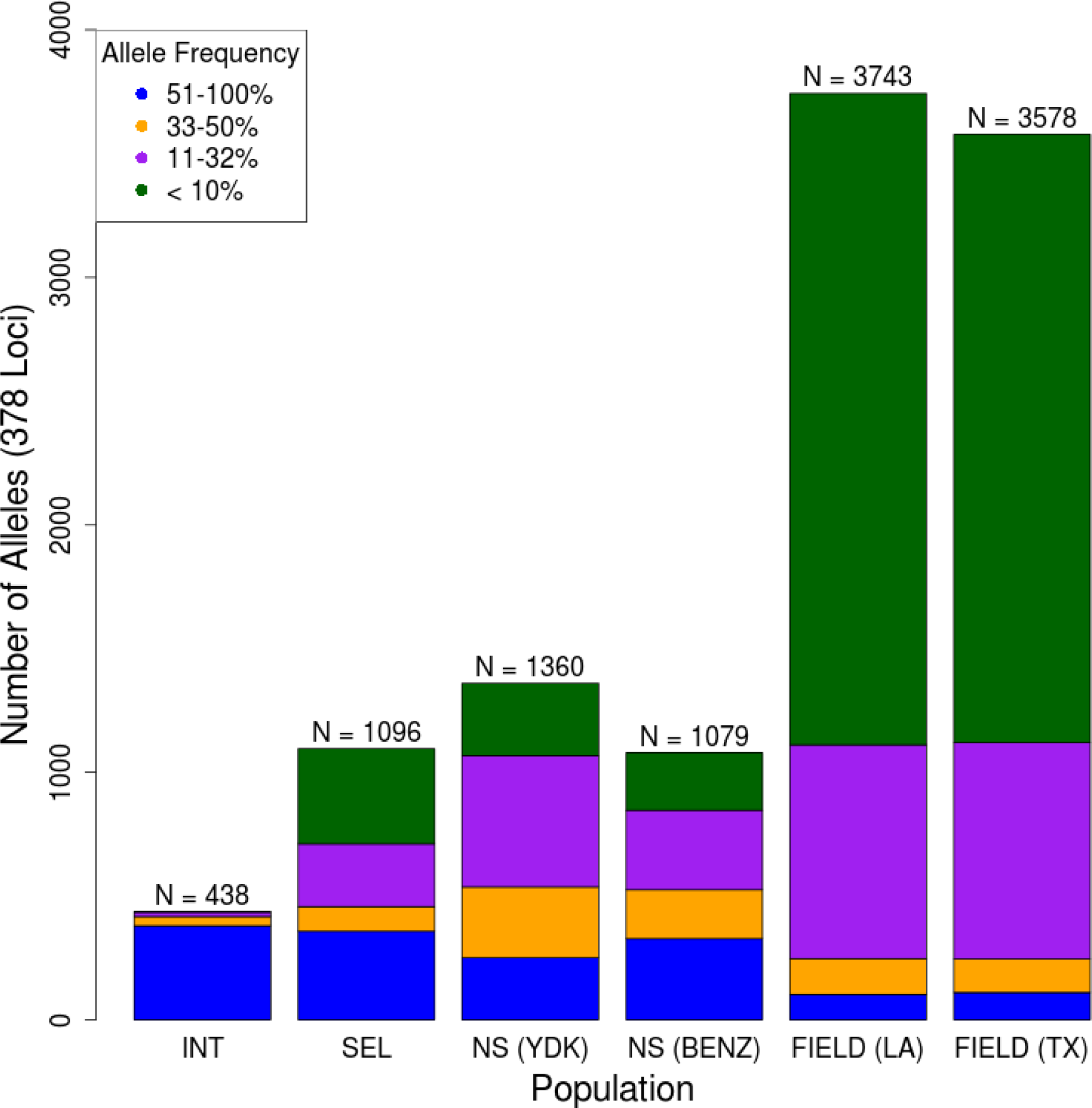
Total numbers of unique alleles detected in sib-mated (INT), Bt-selected (SEL), non-selected (NS), and field-collected (FIELD) populations based upon random sampling of 18 haplotypes per population per locus. Alleles were binned and color-coded according to the frequencies at which they were present out of 18 total haplotypes. Numbers above each bar represent the total number of unique alleles found per population out of 378 loci.

We used the same conservative subset of 378 loci, and calculated sample-size corrected S_K_ (Charlesworth & Charlesworth, 2010), and π (Nei 1978). These two measures are complementary: π is calculated as the proportion of nucleotides that differ per two randomly chosen DNA sequences, averaged across all pairwise comparisons per marker per population, and S_K_ is calculated as the number of unique single nucleotide variants in a population at a single locus. When averaged across all markers (n = 378), the number of variant sites (S_K_) per 350 bp marker was 0.15 for the inbred line, and the maximum S_K_ was 2.95. Bt-selected and non-selected laboratory populations had, on average, just over 1 nucleotide variant per 350 bp locus, with a maximum of *ca*. 6. Field-collected populations had the greatest number of variant sites per 350 bp locus, where the genome-wide average was just over 5 nucleotide variants per locus, with a maximum of *ca*. 15. Similar trends were observed for genome-wide and maximum nucleotide diversity (π) values. Relative to the laboratory-reared populations, genome-wide estimates of π were nearly an order of magnitude lower for the inbred line. The genome-wide π estimate for laboratory-reared populations ranged from 4.0×10^−3^ (Bt-selected population) to 6.7×10^−3^ (non-selected, YDK population), and 6.2×10^−4^ for the inbred line. Field-collected populations exhibited genome-wide π estimates of 9.4×10^−3^ and 9.2×10^−3^ for the LA and TX populations, respectively. Genome-wide and maximum π and S_K_ estimates, along with their corresponding 95% non-parametric bootstrapped confidence intervals (N = 5000) are reported in Table 2.

To further quantify and compare genetic diversity by population, we also examined mean observed heterozygosity in the total population (n = 192 total *H. virescens*), as well as within each sub-population. Observed heterozygosity for the total population (n =192) was 0.27 when averaged across the 378 loci, and considerable variation in heterozygosity existed between populations. Mean observed heterozygosities ranged from 0.06 in the inbred line to 0.40-0.46 in field-collected populations. For laboratory-reared populations, mean observed heterozygosity estimates were intermediate to those of the inbred line and field-collected populations, and ranged from 0.15 in the Bt-selected population to 0.22-0.25 in the non-selected laboratory populations (Table 2).

**Table 2.** Genome-wide nucleotide diversity values per 350 bp locus across populations. All values were calculated using a conservatively chosen set of 378 loci for which at least 10 individuals were genotyped within each population. Genome-wide values represent population-level π and S_K_ averaged across all loci. The abbreviations Bt-sel and NS stand for Bt-selected and non-selected, respectively.

| Population | Mean Observed Heterozygosity | Genome-wide $\pi$ (2.5, 97.5% CIs) | Max $\pi$ | Genome-wide $S_K$ (2.5, 97.5% CIs) | Max $S_K$ |
| --- | --- | --- | --- | --- | --- |
| Inbred line | 0.06 | 0.0006 (0.0004, 0.0008) | 0.013 | 0.15 (0.10, 0.19) | 2.95 |
| Bt-Sel (YHD2) | 0.15 | 0.0040 (0.0035, 0.0043) | 0.020 | 1.67 (1.54, 1.81) | 6.53 |
| NS (YDK) | 0.26 | 0.0067 (0.0061, 0.0073) | 0.028 | 1.66 (1.53, 1.80) | 6.20 |
| NS (BENZ) | 0.22 | 0.0051 (0.0046, 0.0056) | 0.026 | 1.35 (1.24, 1.47) | 5.46 |
| Field (LA) | 0.46 | 0.0094 (0.0087, 0.0100) | 0.034 | 5.35 (5.03, 5.68) | 15.55 |
| Field (TX) | 0.40 | 0.0092 (0.0085, 0.0099) | 0.033 | 5.02 (4.71, 5.35) | 15.63 |

### Genomic divergence among populations

To examine the degree to which the above genetic diversity could be attributed to between population differences, we calculated pairwise estimates of F_ST_ (Table 3) according to Weir and Cockerham (1984). Despite the *ca*. 400 km distance between collection locations for the field populations, very little (0.4%) of the genetic diversity observed in these populations could be attributed to differences between populations. When field populations were compared with non-selected laboratory-reared populations, 16-25% of the genetic variation could be attributed to differences between populations. Additional inbreeding and selection further exacerbated these differences, increasing the percentage of variation attributable to between population differences to over 30%. Of particular interest was the comparison between the ancestral non-selected YDK population, with the more derived Bt-selected (YHD2) and inbred populations. Despite their shared ancestry, a comparison of YDK to YHD2 and the inbred line revealed that 28% and 33% of the existing genetic variation could be attributed to between population differences, respectively.

**Table 3.** Pairwise estimates of genetic divergence across 378 ddRAD-seq loci calculated according to Weir and Cockerham’s *F_ST_*. Estimates of *F_ST_* are above the diagonal, while corresponding bootstrapped confidence intervals (2.5%, 97.5%) are presented below.

|  | **Inbred Line** | **Bt-Sel (YHD2)** | **NS (YDK)** | **NS (BENZ)** | **Field (LA)** | **Field (TX)** |
| --- | --- | --- | --- | --- | --- | --- |
| **Inbred Line** | - | 0.6378 | 0.3272 | 0.6209 | 0.3873 | 0.3899 |
| **Bt-Sel (YHD2)** | (0.5479, 0.7342) | - | 0.2818 | 0.4854 | 0.3137 | 0.3175 |
| **NS (YDK)** | (0.3096, 0.3465) | (0.1823, 0.3796) | - | 0.3863 | 0.1656 | 0.1664 |
| **NS (BENZ)** | (0.5705, 0.6645) | (0.3643, 0.5922) | (0.3345, 0.4288) | - | 0.2552 | 0.2565 |
| **Field (LA)** | (0.3663, 0.4078) | (0.2380, 0.3930) | (0.1521, 0.1850) | (0.2134, 0.2931) | - | 0.0004 |
| **Field (TX)** | (0.3681, 0.4106) | (0.2403, 0.3971) | (0.1529, 0.1861) | (0.2148, 0.2951) | (-0.0115, 0.0156) | - |

### Inbreeding among laboratory-reared *H. virescens* populations

To determine whether the observed degree of heterozygosity in our inbred line was consistent with that which would be expected following 10 generations of sib-mating, we compared the inbreeding coefficient *F*, as calculated according to pedigree-(Falconer & Mackay, 1996) and DNA marker-based information (Keller & Waller, 2002; Kim *et al.*, 2007). The expected inbreeding coefficient (*F_t_*), following 10 generations of sib-mating was 0.89. This expected value fell within the bootstrapped 95% confidence intervals for marker-based inbreeding coefficients (*F_IT_*) calculated from all subsets of markers. This indicated that there was no significant difference between the expected inbreeding coefficient and the observed inbreeding coefficient calculated using ddRAD-seq marker data. The genome-wide *F_IT_* values (95% CIs) were 0.92 (0.88, 0.96), 0.89 (0.86, 0.92), 0.89 (0.87, 0.92), and 0.88 (0.86, 0.89), for the inbred line as calculated from 125, 378, 573, and 1231 ddRAD-seq loci, respectively.

### Linkage mapping

Few genomic resources are available for *H. virescens*. Therefore, we determined the genomic location of loci which were resistant to fixation by generating a dense linkage map. The map was produced via ddRAD-sequencing of the parents and progeny from a male informative cross (reviewed in Baxter *et al.*, 2009). We generated an average of 381,096 sequencing reads (*s.d*. 165,334) per progeny, as well as 493,537 and 762,171 reads per the male and female parents, respectively (Supplementary Figure 3). From this, we produced a linkage map comprised of 659 informative ddRAD-seq loci, plus 3 partial gene sequences of *ABCC2, HevCaLP*, and *DesatI*. Adding these partial gene sequences to our linkage map, all with known locations in the *B. mori* genome allowed us to validate marker groupings for our linkage map. All informative ddRAD-seq loci were grouped into 33 linkage groups, two more than the expected 30 *H. virescens* autosomes, and one segregating Z chromosome from the hybrid male parent used in our cross. Linkage groups ranged in size from 7cM to 110cM (Figure 3), and yielded a total map length of 1919.5 cM. On average, there were 20 ddRAD-seq loci per linkage group, and the average spacing was one locus per 3.5 cM. The smallest and largest linkage groups contained 3 and 53 loci, respectively. The *HevCaLP*, *Desat1*, and *ABCC2* genes were grouped with linkage groups 15, 16, and 22, respectively. These linkage groups corresponded to *B. mori* chromosomes 6, 23, and 15 (Table 4), where these candidate genes are known to reside (Gahan *et al.*, 2001; Gahan *et al.*, 2010; Mita *et al.*, 2004).

**Figure 3.**
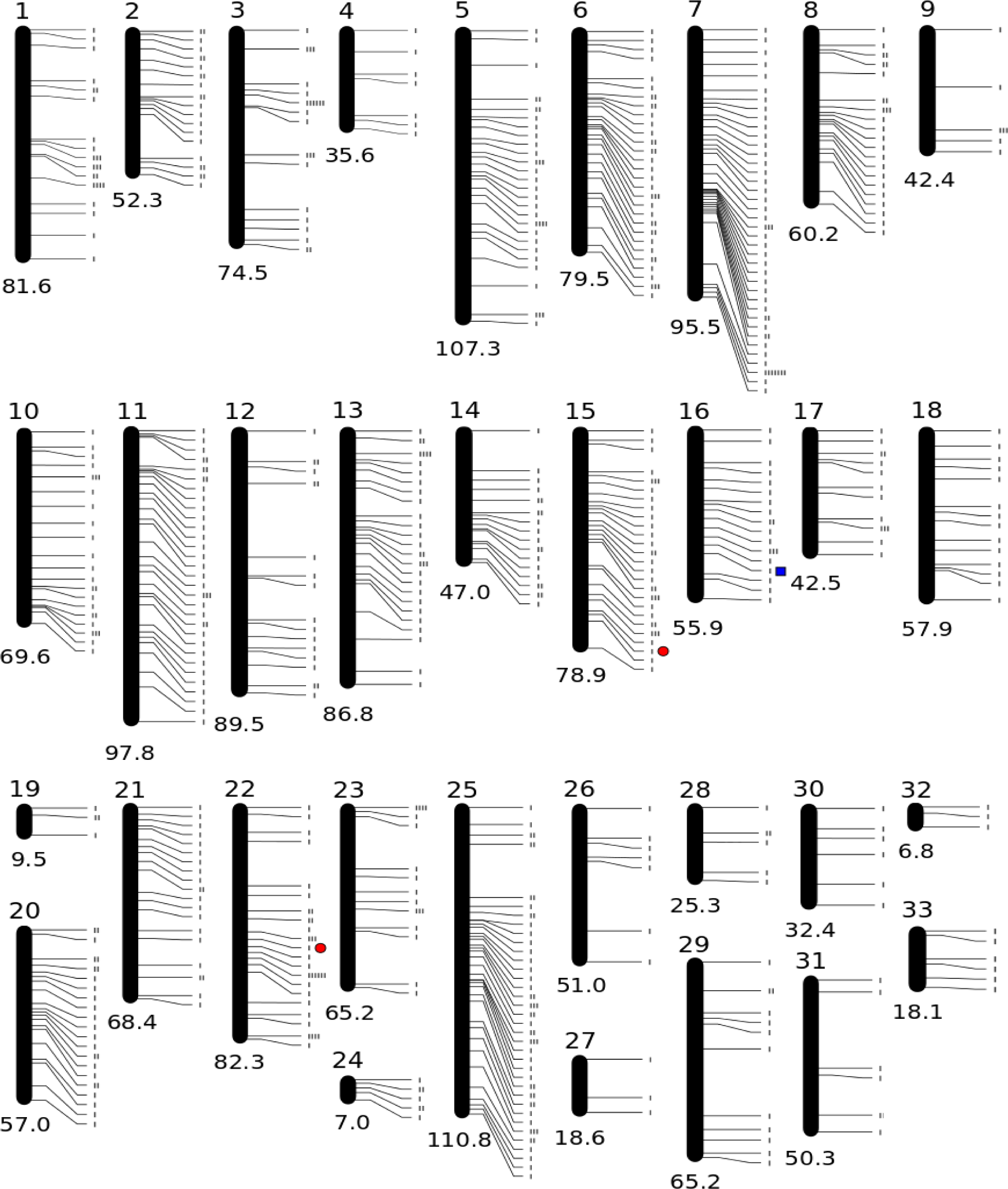
*Heliothis virescens* linkage map with a total length of 1919.5 cM. Centimorgan lengths are below each linkage group. Each tick mark represents an individual marker that mapped to a particular position in the linkage group. Red circles next to linkage groups 15 and 22 represent the positions of the *HevCaLP*, and the *ABCC2*, respectively. The blue square represents the position of the delta-11-desaturase on linkage group 16.

**Table 4.** Linkage group (LG) correspondence with *B. mori* chromosome (Chr). Linkage groups with an asterisk contained one or more markers that aligned uniquely to an unmapped *B. mori* sequence. Where linkage groups contained markers that aligned to more than one *B. mori* chromosome, italicized marker names correspond to the italicized *B. mori* chromosome.

| LG | *B.mori* Chr | Number Markers<br>Aligned to *B.mori*<br>Chr | Total<br>markers in<br>LG | LG Length<br>(cM) | Average Marker<br>Spacing (cM) | Names of Markers Aligned to *B.mori* Chr |
| --- | --- | --- | --- | --- | --- | --- |
| 1 | 25 | 3 | 25 | 81.6 | 3.3 | 19754, 29329, 20282 |
| 2 | 8 | 6 | 23 | 52.3 | 2.3 | 22394, 27499, 21475, 17667, 17852, 25500 |
| 3* | 17 | 3 | 24 | 74.5 | 3.1 | 376, 21931, 25112 |
| 4 | - |  | 7 | 35.6 | 5.1 |  |
| 5* | 4 | 5 | 36 | 107.3 | 3.0 | 2556, 29595, 22095, 8695, 15160 |
| 6 | 5 | 6 | 40 | 79.5 | 2.0 | 18575, 4268, 21041, 23679, 66, 5723 |
| 7 | 22 | 9 | 49 | 95.5 | 1.9 | 13710, 2328, 1443, 3654, 13880, 22897, 3123,<br>2430, 19858 |
| 8* | 11 | 9 | 25 | 60.2 | 1.7 | 22584, 939, 3060, 18820, 23156, 17932, 202,<br>22290, 21173 |
| 9 | 23 | 2 | 7 | 42.4 | 6.1 | 4275, 1286 |
| 10 | 3 | 5 | 26 | 69.6 | 2.7 | 5015, 5329, 4265, 4545, 23262 |
| 11 | 9 | 8 | 37 | 97.8 | 2.6 | 19938, 6113, 16969, 19392, 1572, 89,<br>22588, 19343 |
| 12 | - |  | 18 | 89.5 | 5.0 |  |
| 13 | 13 | 6 | 31 | 86.8 | 2.8 | 2074, 15962, 26388, 1193, 16684, 18825 |
| 14 | 19 | 1 | 21 | 47.0 | 2.2 | 14242 |
| 15 | 6 | 7 | 35 | 78.9 | 2.3 | 25943, 4579, 25083, 1752, 3385, 2784, 18178 |
| 16 | 23 | 4 | 20 | 55.9 | 2.8 | 2485, 16350, 806, 20599 |
| 17 | 7 | 2 | 14 | 42.5 | 3.0 | 29343, 343 |
| 18 | 26 | 1 | 14 | 57.9 | 4.1 | 1113 |
| 19 | 11, 22 | 2 | 4 | 9.5 | 2.4 | 3536, *16185* |
| 20* | 10 | 6 | 25 | 57.0 | 2.3 | 17820, 2978, 18970, 12277, 16316, 12984 |

**Table 4 continued.**
| LG | *B.mori* Chr | Number Markers<br>Aligned to *B.mori*<br>Chr | Total<br>markers in<br>LG | LG Length<br>(cM) | Average Marker<br>Spacing (cM) | Names of Markers Aligned to *B.mori* Chr |
| --- | --- | --- | --- | --- | --- | --- |
| 21 | 21 | 3 | 21 | 68.4 | 3.3 | 26113, 20542, 7624 |
| 22 | 15 | 2 | 33 | 82.3 | 2.5 | 21411, 5773 |
| 23 | 14 | 1 | 17 | 65.2 | 3.8 | 4156 |
| 24 |  |  | 7 | 7.0 | 1.0 |  |
| 25 | 12,1 | 4 | 53 | 110.8 | 2.0 | 20280, 9211,12382, 23604 |
| 26 |  |  | 7 | 51.0 | 7.3 |  |
| 27 | 2 | 1 | 3 | 18.6 | 6.2 | 5781 |
| 28 |  |  | 6 | 25.3 | 4.2 |  |
| 29 | 16 | 1 | 12 | 65.2 | 5.4 | 23481 |
| 30 |  |  | 6 | 32.4 | 5.4 |  |
| 31 |  |  | 7 | 50.3 | 7.2 |  |
| 32 |  |  | 3 | 6.8 | 2.3 |  |
| 33 | 14, 28 | 2 | 6 | 18.1 | 3.0 | 6601, 20184 |

In total, 99 of the 659 mapped ddRAD-seq loci could be aligned uniquely to a single locus in the *B. mori* genome. Twenty-two linkage groups contained ddRAD-seq loci that could be aligned to a single *B. mori* chromosome (Table 4), while 8 linkage groups did not contain any that could be aligned. Linkage groups 19, 25, and 33 contained ddRAD-seq loci that aligned uniquely to more than one *B. mori* chromosome. This was unlikely caused by spurious associations between ddRAD-seq loci; increasing the LOD score to 8 failed to break up associations for those three linkage groups.

### Identification of genomic regions resistant to fixation

To determine where genetic diversity was being maintained in the genome, we examined observed heterozygosity and nucleotide diversity at the 659 mapped ddRAD-seq loci for one field-collected (LA), the Bt-selected (YHD2), one non-selected (YDK), and inbred lines. Of these 659 mapped loci, 302 (46%) were previously included in our population-level analyses, and 357 had not been previously analyzed. For each mapped locus, observed heterozygosity and nucleotide diversity (π) values were only calculated if at least 3 individuals were genotyped per population. Therefore only 441 loci were examined for the inbred line, 658 loci were examined for the non-selected (YDK) population, 659 loci were examined for the Bt-selected population, and 546 loci were examined for the field-collected population. In total, 13% (n = 60) of mapped loci retained polymorphism in the inbred line, whereas 98% (n = 645), 86% (n = 583), and 99% (n = 543) of mapped loci retained polymorphism in the Bt-selected, non-selected (YDK), and field-collected (LA) populations, respectively. For each of these populations, levels of observed heterozygosity and nucleotide diversity across the genome are compared in Figure 4.

**Figure 4.**
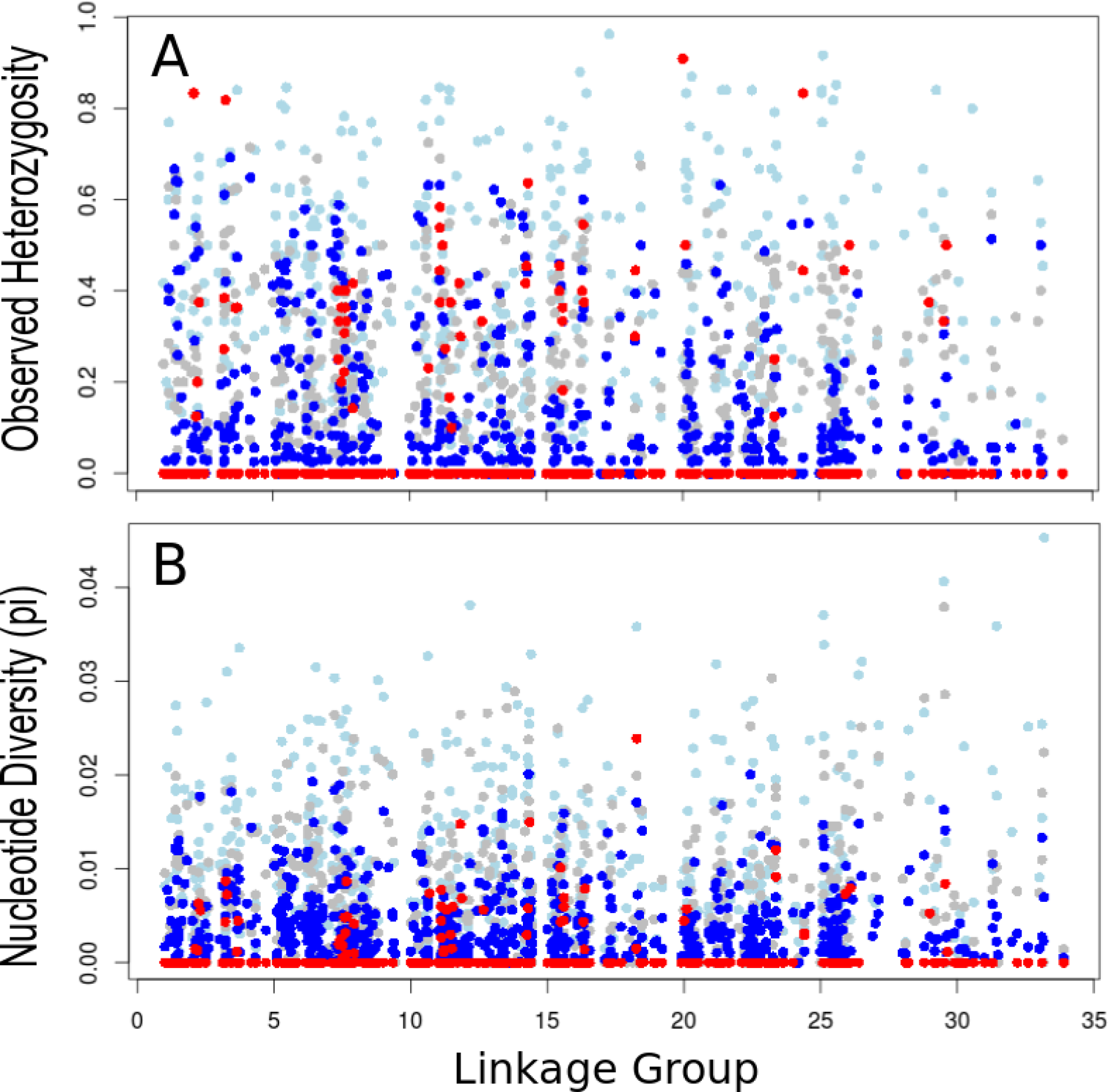
Observed heterozygosity (A) and nucleotide diversity (B) estimates per mapped marker site. For comparative purposes, estimates for the Louisiana field population were included (light blue). Grey and blue circles represent estimates in non-selected (YDK), and Bt-selected (YHD2) populations, respectively. Red circles represent estimates in the inbred line following 10 generations of sib-mating.

Within the inbred line, mapped loci that retained polymorphism were spread over 16 linkage groups (Figure 4), and often appeared in clusters within a linkage group (Supplementary Figure 4). Linkage groups 7 and 11 had particularly high numbers of polymorphic markers relative to other linkage groups. Therefore, we examined whether clustering of polymorphic markers in our inbred line was greater than could be expected due to random chance. Such clustering may point to the presence of large chromosomal inversions as observed in other insect species (Turissini et al. 2014). A replicated G-test of independence (Sokal and Rohlf, 1995) demonstrated that 8 linkage groups contained significantly more polymorphic loci than would be expected following 10 generations of inbreeding (Supplementary Table 2). Yet based upon the non-significant heterogeneity g-value (G = 23.87, df = 23, p-value = 0.41) we could not reject the null hypothesis that the distribution of polymorphic loci was homogeneous across linkage groups, and the clustering observed within some linkage groups did not significantly differ from that which could be observed by chance.

We also examined whether balancing selection, perhaps due to balanced lethal systems, may be responsible for the residual polymorphism observed in the inbred line. To this end, we calculated and compared Tajima’s D values at each polymorphic locus for the inbred line and it’s ancestral population (YDK). Tajima’s D is a statistical test which identifies departures from the neutral model of molecular evolution. A positive Tajima’s D value at a locus indicates an excess of intermediate frequency alleles, thereby signifying either a recent population contraction, or balancing selection at that locus. We reasoned that positive Tajima’s D values in the inbred line would be much less likely to signify a recent population contraction, and therefore more likely to indicate balancing selection, if these same regions were also significant in their ancestral population (YDK). Indeed, YDK had not undergone any obvious population contractions around the time of this separation 10 generations prior. Therefore, we examined which, if any, of these polymorphic loci in the inbred line shared strongly positive Tajima’s D values with YDK. When we calculated Tajima’s D values for the 60 polymorphic sites remaining in the inbred line, 21 were significantly positive (α = 0.05). Following a Benjamini-Hochberg correction for multiple comparisons (Benjamini and Hochberg, 1995), only two loci remained statistically significant. Furthermore, neither of these two polymorphic loci from the inbred line overlapped with loci showing significantly positive Tajima’s D values in YDK. Thus we found little evidence of ongoing balancing selection in our inbred line.

## Discussion

Here, we examined the degree to which colonization, artificial selection, and intense inbreeding influence genome-wide and fine-scale patterns of diversity. In the absence of a publicly available *H. virescens* reference genome, we used ddRAD-seq *de novo* locus construction to identify multiple subsets of polymorphic loci ranging in size from 125 to 1231 markers. Genome-wide measures of allelic diversity, *F_IT_* values, and the degree of homozygosity were either unaffected (Supplementary Figure 2), or minimally affected (Supplementary Table 1) by inclusion of markers with high levels of missing genotypic data. Therefore, any biased genotype calls made by the Stacks SNP calling algorithm due to our moderate depth of sequencing coverage had little impact on our overall genome-wide estimates of diversity. Our results demonstrate that moderate coverage ddRAD-seq data can be used with confidence when conducting population genomic comparisons of genome-wide means.

We observed a precipitous decline in nucleotide and allelic diversity following long-term laboratory colonization, selection, and inbreeding for *H. virescens*. Despite the decline in genomic diversity for non-selected and Bt-selected laboratory-reared populations, fewer than 10% of loci were fixed. While our Bt-selected population did not retain the level of genetic diversity that their ancestral (YDK) laboratory-reared population did, they consistently had higher measures of genomic diversity than did the non-selected (BENZ) population. Retention of higher levels of polymorphism in our Bt-selected line was likely due to the measures taken during its generation to ensure genomic diversity was maintained in the face of strong selection (Gould, 1995). Alternatively, strong genetic bottlenecks in the non-selected (BENZ) population prior to their use in our study could explain why our Bt-selected line was more genetically diverse than the BENZ non-selected line. Overall, differences among Bt-selected and non-selected laboratory-reared populations were modest; when 18 total alleles were sampled, laboratory-reared populations retained *ca*. 3 alleles per 350-bp locus relative to 9 alleles per 350-bp locus present in field-collected *H*. *virescens*. However, few low frequency alleles remained in the laboratory-reared populations relative to the field-collected populations (Figure 2), which has been observed elsewhere (Munstermann, 1994). For this reason, laboratory-reared populations are generally considered inbred (Roush, 1986). In the case of *H. virescens*, our results clearly show that a great deal of genomic diversity is retained, even following decades in colony. Few genome assembly algorithms accommodate polymorphism well (Kajitani *et al.*, 2014), and it is clear that the reductions in heterozygosity in our inbred line will be useful for production of a high quality *H. virescens* reference genome assembly.

To determine the degree to which the genomic diversity detected above could be attributed to differences between populations, we calculated pairwise *F_ST_* values according to Weir and Cockerham (Table 3). An *F_ST_* value of 0.0004 demonstrated that most of the existing genetic diversity occurred within, as opposed to between, these two field-collected populations of *H. virescens*. This result is similar to that of Groot *et al*. (2011), which described over 98% of the genetic variation detected across North American *H. virescens* to be found within populations. Both studies suggest that extensive gene flow occurs naturally among geographically disparate *H. virescens* populations. Although we did not sample field-collected populations from Yadkin County, NC, nor from Stoneville, MS, the original collection sites of our laboratory-adapted YDK and BENZ populations, our results along with those of Groot et al. (2011), suggest that genomic divergence among all four sites would have been low. To examine how decades of laboratory rearing could influence the structure of genomic diversity in laboratory-adapted populations, we compared our YDK and BENZ populations to our field-collected populations. As would be expected, we observed an increase (16-25%) in the percentage of genetic variation that could be attributed to between population differences. This is likely due to the purging of rare alleles (as shown in Figure 2) and the fixation of others as caused by the random process of genetic drift. Further manipulations, like Bt-selection and full-sibling mating increased that percentage of between population genetic variation to well over 30% relative to the field-collected populations. Additional comparisons between ancestral YDK and the more derived Bt-selected and inbred populations further underscore the affects that these population-level manipulations can have on insect colonies in a laboratory setting. Although they were derived from YDK, following Bt-selection and over 2 decades of separation in the laboratory, 28% of the total genetic variability could be attributable to between-population differences. Likewise, 10 generations of full-sibling mating resulted in over 32% of the total genetic variability existing between YDK and the inbred line.

Following 10 generations of inbreeding, over 80% of markers went to fixation in our sib-mated *H. virescens* population. Indeed, our inbreeding coefficient *F_IT_*, as observed from our ddRAD-seq data, met theoretical expectations for all subsets of loci. Our *H. virescens* laboratory population was more amenable to inbreeding than other insect species (Munstermann, 1994; Rumball *et al.*, 1994; Turissini *et al.*, 2014; You *et al.*, 2013), despite their relatively high levels of genomic diversity (Figure 1) and genetic load (Supplementary Figure 5). Higher than expected allelic diversity has been observed in several other insect species following experimental inbreeding attempts (Munstermann, 1994; Rumball *et al.*, 1994; Turissini *et al.*, 2014; You *et al.*, 2013). As one example, only 57% of the *An. gambiae* genome went to fixation, as observed according to SNP markers, following 10 generations of inbreeding (Turissini *et al.*, 2014). Observed differences between our *H. virescens* population and other insects could be species specific, but is more likely related to the proportion of the genome containing balanced lethal systems (Falconer & Mackay, 1996).

To determine where heterozygosity was being maintained in our inbred line, we developed a high density genetic linkage map for *H. virescens*. Our map contained 659 newly developed 350 bp ddRAD-seq markers that are long enough for future primer design and direct sequencing. This map represents a new tool for an historically important pest species that lacks genomic resources. No ddRAD-seq markers remained unlinked following mapping, which indirectly speaks to the quality of our linkage map. However, the number of groups in our linkage map was 2 more than expected (n = 31). This is likely due to the relatively small mapping population size used in this work; other mapping studies that analyzed segregating populations of a similar size have also reported genetic maps with excess numbers of linkage groups (Pootakham *et al.*, 2015; Singh *et al.*, 2009). Additional explanations for the disparity between our observed and expected number of linkage groups include the uneven distribution of markers over the chromosomes (Paterson, 1996), or recombination ‘hotspots’, which make it difficult to reduce the number of linkage groups to 31.

When we applied our linkage map to examine fine-scale patterns of genomic change following 10 generations of sib-mating, we found that several linkage groups seemed to contain clusters of loci that retained polymorphism (Figure 4). However, results from a replicated G-test of independence demonstrated that clustering of these polymorphic loci was not significantly different from what could be expected due to chance. This does not preclude the possibility that chromosomal inversions exist among populations of *H. virescens*. Rather it suggests that there were no large inversions responsible for maintaining polymorphisms in our inbred line.

Furthermore, only 2 polymorphic loci had significantly positive Tajima’s D values, and neither overlapped with those loci that had high D values in YDK. This made it impossible to rule out population contraction as a contributor to the positive Tajima’s D values. Unlike the findings for other insects (Mackay et al. 2012, Turissini et al. 2014), the combined results of our *F_IT_* test, replicated G-test, and Tajima’s D tests suggest that neither balanced lethals nor large chromosomal inversions appear to play a major role in retention of polymorphism for this particular *H. virescens* inbred line.

## Conclusions

This work serves as one of the most thorough attempts to quantify the effects of genomic responses to selection and inbreeding in a non-model insect species. We demonstrated that laboratory-reared *H*. *virescens* have reduced allelic and nucleotide diversity relative to field-collected populations, and that inbreeding further diminishes genetic diversity. Although we identified several loci that did not go to fixation in *H. virescens* following 10 generations of inbreeding, our ddRAD-seq marker-based *F_IT_* values met theoretical expectations. This work demonstrates the difficulty involved in producing fully homozygous insect strains, which are currently critical to producing high-quality, complete reference genomes.

## Methods

### *Field-collected* H. virescens

Adult male moths were collected from Bossier Parish, Louisiana, and College Station, Texas using pheromone-baited live traps. Collections took place in LA from May through September, 2012, and in TX from May through October, 2012. Moths from each collection date were immediately placed in bottles of 95% ethanol for long-term storage. All bottles were held at -20 °C until DNA isolations took place.

### H.virescens *colonies*

*H.virescens* were collected from Yadkin County, NC in 1988 (Gould *et al.*, 1995). This original population founded two of the colonies used in this study, each of which had been reared in the laboratory for *ca*. 290 generations. YHD2 was selected for high levels of Bt resistance for 4 years (up to 48 generations) on MVP-treated (0.864 mg/mL diet; Mycogen, San Diego, CA) corn-soy diet (Gould *et al.*, 1995), whereas a non-selected population (YDK) was reared on corn-soy diet alone. A third population (BENZ) originating from Stoneville, MS, was acquired from Benzon Research Incorporated (Carlisle, PA) and had been reared in the laboratory for over 10 years (120 generations). BENZ *H. virescens* were acquired in their pupal stage, and newly eclosed adults were used for population-level comparisons. To produce an inbred population, single pair matings (SPMs) were set up between YDK siblings for 10 generations. An initial 37 SPMs were used to establish 29 lineages in filial generation one (8 single pair matings did not produce progeny). When SPMs failed to produce offspring, likely due to inbreeding depression, surviving lineages were expanded (Supplementary Figure 5). This was done to extend inbreeding for as many generations as possible, thus promoting as complete a reduction in heterozygosity as possible. Adult males from each laboratory-reared population were killed by freezing (-20 °C), and stored at -80 °C until DNA isolation.

### Mapping cross

A non-selected female from the BENZ population was crossed to a Bt-selected (YHD2) male in a single pair mating. One hybrid male offspring was then back-crossed to a Bt-selected (YHD2) female, and their progeny were reared to adulthood on untreated corn-soy diet according to Joyner and Gould (1985). Of the 120 progeny, 97 reached adulthood. Parents and their 97 adult progeny were killed by freezing and stored until DNA isolation as described above.

### Genomic DNA library preparation

All DNA was isolated from the adult thorax using a Qiagen Dneasy Blood and Tissue Kit (Qiagen, Inc., Valencia, CA, U.S.A.). Genomic DNA samples were prepared for Illumina sequencing according to the Poland *et al.*, (2012) protocol with minor modifications. Two-hundred ng of DNA per individual were digested with EcoRI and MspI. For each individual, the overhang sites were ligated to standard Truseq Universal adapters (Illumina, Inc. San Diego, CA). Adapters ligated to EcoRI overhang sites contained one of 48 unique barcodes (Elshire *et al.*, 2011; Supplementary Table 3). DNA fragments from each individual were assigned a unique barcode, and individuals were combined into pools of no more than 48 individuals. A Pippin Prep (Sage Science, Inc., Beverly, MA) was used to select adapter-ligated DNA fragments ranging from 450-650 bp from each pool. Size-selected DNA fragments were amplified in a Peltier PTC200 thermalcycler (here and throughout) using Illumina primers (Supplementary Table 4) under the following conditions: 72 °C for 5min, 18 cycles of 98 °C for 30 sec, 65 °C for 20 sec, 72 °C for 30 sec, followed by 72 °C for 5 min. For each pool, 1 of 4 Illumina indices was added via PCR to the MspI adapter. Therefore, sequences from each individual could be identified by the unique combination of barcode and index. A complete list of barcodes and indices used in this study can be found in the Supplementary Tables 3 and 4, respectively. Amplified libraries were pooled, cleaned with a Qiaquick PCR Purification Kit (Qiagen, Inc., Valencia, CA, U.S.A.), and diluted to 4nM prior to sequencing. Prepared genomic DNA libraries constructed from 303 *H. virescens* individuals were spread across 9 full and partial Illumina MiSeq runs. The MiSeq reagent kit v. 2 was used for initial preparation of the mapping family and the inbred line. All subsequent preparations, including re-runs of the mapping family and inbred line were prepared with the MiSeq reagent kit v. 3.

### De novo *marker formation*

Overlapping paired-end reads were merged with FLASH (Magoc & Salzberg, 2011), and Stacks v. 1.09 (Catchen *et al.*, 2011; 2013) was used for demultiplexing and *de novo* formation of loci. Merged paired-end reads were filtered for quality using the process_radtags script. Further quality filtering entailed removal of reads when: 1) they did not have an intact EcoRI cut site, 2) had a quality score < 30, or 3) were smaller than 350 bp. We did not allow process_radtags to rescue reads where barcode sequences contained an error. All remaining merged reads were truncated at a length of 350 bp, and fed into the Stacks pipeline.

### Stacks parameter settings

Reads from all individuals were run through ustacks with the following parameter settings: -m 3, -M 14 (allowing for 5% nucleotide mismatch rate between alleles per individual), -max_locus_stacks 2, —alpha 0.05. A consensus catalog of loci was first formed using the parents of the mapping cross with cstacks, where the -n 14 parameter allowed for a 5% between individual nucleotide mismatch rate. For the mapping family, genotype calls were made using sstacks prior to field- and colony-strain alleles being added to the catalog. Progeny genotypes were automatically corrected using the Stacks genotypes script. Twenty-four individuals of each colony and field-collected strain collected in 2012 were later added to the catalog, and all field-collected and laboratory-reared populations were also genotyped using sstacks.

### Data analyses

All population genomic and linkage analysis were conducted in R version 3.1.2 (R core team, 2014).

### *Genomic diversity among* H. virescens *populations*

In total, we sequenced the 13 surviving males from an inbred line subjected to 10 generations of sib-mating, 42-46 males per colony strain, and 30 males per field-collected population (Table 1). Prior to running sequence data through the Stacks pipeline, we checked individual read counts across populations to ensure uniformity (Supplementary Figure 1). Twelve of the 204 individuals sequenced had too few (< 90,000) or too many reads (> 688,000) and were removed from the dataset prior to analysis, following Bi *et al*. (2013). From our Stacks output, we constructed 4 different sets of consensus loci present across populations. These subsets, containing a core overlapping set of 125 loci, and increasing in size from 125 to 1231, consisted of marker sets with varying percentages of missing genotype calls (range = 11.2-29.5%) (Supplementary Table 1). We used these 4 different subsets to examine and compare changes in genomic diversity across populations.

We estimated the mean number of unique alleles present per locus, and corresponding 95% non-parametric bootstrapped confidence intervals (N = 5000) across populations using a custom-written R script. Each allele represented the accumulation of SNPs within a 350bp locus, analogous to a haplotype. At a given locus, and within each population of size N (see Table 1 for sample sizes), alleles (total = 2N) were randomly sampled without replacement either 6, 12, 18, or 24 times. Then the number of unique alleles were counted for each sampling regime. Due to their small sample size resulting from intensive inbreeding, we only sampled 6, 12, and 18 alleles per locus for the inbred line. Our analysis focused primarily on the subset of loci containing 378 consensus loci because genotype calls were present for at least 10 individuals per population. We also calculated two measures of nucleotide diversity per 350 bp locus using the R package, pegas (v. 0.6; Paradis, 2010): π (Nei, 1987) and S_K_ corrected for sample size (Charlesworth & Charlesworth, 2010; Watterson, 1975). We then generated population-level genome-wide means and 95% non-parametric bootstrapped (N = 5000) confidence intervals for each metric (Table 2).

### Genetic divergence between populations

To determine the degree of genetic diversity accounted for by differences between our field-collected, laboratory-reared, Bt-selected, and inbred populations, we calculated Weir and Cockerham’s *F_ST_* (Weir and Cockerham, 1984) along with corresponding 95% bootstrapped confidence intervals (N = 5000). Calculations were carried out using the R package, diveRsity (v. 1.9.89; Keenan *et al.*, 2013).

### Estimating the inbreeding coefficient

To estimate our marker-based inbreeding coefficient, we examined multiple sets of loci (Supplementary Table 1) and found that trends across all datasets were similar (data not shown). However, we reported *F_IT_* values from a set of 378 loci because this reduced dataset contained few missing genotypes per population, while still making inferences from several hundred loci. We calculated *F_IT_* for the inbred line relative to the non-selected (YDK) population after Keller and Waller (2002), where 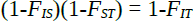. *F_IS_* was the level of inbreeding within the inbred line, calculated as 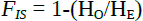, where H_O_ and H_E_ were calculated for each locus using the R package adegenet (v. 1.4-2; Jombart, 2008). *F_ST_* was the accumulated effect of inbreeding over time, calculated as 1-(H_E_(inbred line)/H_E_(YDK)).

### Amplification and genotyping of PCR-based markers for the mapping family

Progeny from the mapping family were genotyped at three additional loci, and these loci were mapped alongside our ddRAD-seq markers to validate our linkage groupings. We targeted the previously described *H. virescens* genes *DesatI, ABCC2*, and *HevCaLP* via PCR followed by gel electrophoresis, or direct sequencing. Amplification and genotyping protocols were as follows.

A 468 bp fragment from *DesatI* was amplified in a 30 μl reaction with forward and reverse primers [5’-TGAGGGACCATCGTCTCCAT-3’] and [5’-CACTGCTACATTTTGGGCAG-3’], respectively (Ward, 2009). Each reaction contained 6 μL of 5× GoTaq buffer (ProMega), 29 μM per dNTP, 92 ng per primer, 0.75 U GoTaq polymerase, and *ca*. 1 μg genomic DNA. Sample DNA was amplified alongside a negative control (here and throughout), where pcr-grade H_2_O was substituted for genomic DNA. Reactions were incubated at 95°C for 1 min followed by 35 cycles of 95°C for 1min, 52°C for 1min, and 72°C for 2 min. PCR products were purified using a standard ethanol precipitation, and directly sequenced on an ABI3730xl (Applied Biosystems, San Francisco, CA). A single nucleotide polymorphism (cytosine to tyrosine substitution) at bp 36 was found in the YHD2 parent of the mapping cross. Offspring were genotyped at this locus using PolyPhred (Nickerson *et al.*, 1997), and genotype calls were visually confirmed using consed (Gordon *et al.*, 1998).

An intronic region of the *ABCC2* gene previously described by Gahan *et al*. (2010) was amplified using primers Hs-ABC2dU02-F1 [5’ – TGGTTACAAGAAATAGAAAATGCAAC – 3’] and Hs-ABC2eU03-R2 [5’ – CTTTCAAACTGAACCGCATCAC – ’3]. Each 30 μl reaction volume consisted of 6 μL of 5× GoTaq buffer, 29 μM per dNTP, 73 ng per primer, 0.75 U GoTaq polymerase, and 1 μg genomic DNA. Reactions were held at 95°C for 2 min followed by 30 cycles of 95°C for 30 sec, 58°C for 30 sec, and 72°C for 40 sec, and the resulting products were cleaned via ethanol precipitation. Following sequencing on an ABI3730xl, chromatogram files were visualized using FinchTV (version 1.3.1, PerkinElmer, Inc., Seattle, WA). As described by Gahan *et al*. (2010), the YHD2 parent was homozygous for a 22 bp deletion, whereas the F1 parent was heterozygous for this deletion. Therefore, we examined the segregation of this deletion in the mapping family offspring, which was detectable by the presence of a TAT sequence near amplicon bp 40.

Finally, the *HevCaLP* locus described by Gahan *et al*. (2007) was amplified in a multiplexed reaction using three primers: the universal reverse primer [5’ – ATACGAGCTGACGACACGCTGGGAGA – 3’], one forward primer that targets a retrotransposon insertion conferring resistance to *Bacillus thuringiensis* [5’ – CGCAACGCGCGATCTACTCTTGTCACC – 3’], and another forward primer that targets wild-type sequence [5’ – AAGTGTCCCAGTCGATGCTGAA – 3’]. An initial 20-μl reaction contained 4 μl 5× GoTaq buffer, 29 μM per dNTP, 56 ng per primer, 0.5 U GoTaq polymerase, and 1 μg genomic DNA. Reactions were incubated at 95°C for 2 min, followed by 30 cycles of 95°C for 30 sec, 58°C for 20 sec, and 72°C for 40 sec. A reconditioning reaction, aimed at reducing heteroduplex formation, was set up as above, but incubated for 3 rather than 30 cycles. These reactions were capable of producing two amplicons, which differed in length by 76 bp. The YHD2 parent was homozygous for the long amplicon (*ca*. 800 bp) containing the insertion that confers resistance to *Bacillus thuringiensis*, whereas the F1 parent was heterozygous for a long and short amplicon. PCR products from mapping family offspring were run on a 2% agarose gel alongside Hyperladder I (Bioline, Taunton, MA) for visualization and genotype scoring.

### Linkage mapping

Double-digest RAD-seq markers present in fewer than 75% of the mapping family offspring were filtered out, and the remainder were checked using a chi-square test for Mendelian segregation (α = 0.01). PCR-based markers, as well as those ddRAD-seq markers that segregated in a mendelian fashion were assigned to linkage groups (LOD = 5, maximum recombination fraction = 0.3) using the onemap package (Margarido *et al.*, 2007) in R. We validated groupings by aligning all markers to the *Bombyx mori* genome using Blastn in Kaikobase version 3.2.2 (http://sgp.dna.affrc.go.jp/KAIKObase/). Furthermore, we confirmed that the locations of the *DesatI, ABCC2*, and *HevCaLP* pcr-based markers, as well as ddRAD-seq markers found in their respective linkage groups aligned to the same *B. mori* chromosomes (Table 4). Markers on each linkage group were ordered using the recombination counting and ordering algorithm (RECORD; Van Os *et al.*, 2005). RECORD was chosen based upon previous studies demonstrating the reliability of its performance (Collard *et al.*, 2009, Mollinari *et al.*, 2009). Recombination fractions were converted to centiMorgan distances using the Kosambi mapping function (Kosambi, 1944). The final linkage map was drawn using Genetic Mapper version 0.5.

### Assessment of fine-scale differences in nucleotide diversity across laboratory-reared populations

Mapped markers were examined for observed heterozygosity (H_o_) and nucleotide diversity (π), as above, for the inbred line, the Bt-selected line, one non-selected line (YDK), and one field-collected population (LA), and these values were displayed for visual comparison in Figure 4. For each population, at least 3 individuals must have been genotyped for a marker to be included in the analysis. Then, we examined the distribution of mapped markers that retained polymorphism in the inbred line to determine whether heterogeneity, or significant clustering, could be observed. Using a replicated G-test of independence (Sokal and Rohlf, 1995), the distribution of polymorphic loci across all linkage groups that contained five or more markers (24 of the 33 total linkage groups) was examined. Under assumptions of homogeneity, we expected that the ratio of polymorphic to fixed loci across linkage groups in the inbred line would follow Hartl and Clark’s (2007): 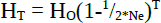, where H_O_ = initial observed heterozygosity in the population, which we set equal to the observed heterozygosity of it’s ancestral YDK population.

Ne = the number of breeding adults in each generation, which we set equal to 2 according to our sib-mating design.

T = the number of generations

According to this equation, the expected frequency of polymorphic loci was 0.015 within a linkage group following 10 generations of sib-mating. Each linkage group was first examined for deviation from this expected frequency of polymorphic loci using a G-test of independence with a Bonferroni-adjusted alpha value (α = 0.002) to account for multiple comparisons. Individual uncorrected G-values produced for each of the 24 linkage groups were added together to generate a “pooled” G-value, and a “total” G-value was calculated according to Sokal and Rohlf (1995). Finally, a heterogeneity G-value, which tested the hypothesis that polymorphic loci were significantly more clustered within linkage groups than due would be due to random chance, was calculated by taking the difference between total and pooled G-values, then comparing it to a Χ^2^ distribution with 23 degrees of freedom and an apriori alpha value of 0.05.

Each marker that retained polymorphism in the inbred line was then examined for an excess of intermediate frequency alleles, which could indicate either a population contraction or ongoing balancing selection near that locus. For this, we used a Tajima’s D test as calculated by the R package, pegas (v. 0.6; Paradis, 2010). To tease apart the effects of population demographic changes and forces of natural selection, we compared the Tajima’s D values at each locus in the inbred line to those of their ancestral population (YDK). We reasoned that significantly positive Tajima’s D values that overlapped between the two lines would be less likely due to the population demographic changes created by sib-mating, and more likely due to selection. A Benjamini-Hochberg adjustment was applied to all Tajima’s D p-values to account for multiple hypothesis tests using the R package fdrtool (v.1.2.15; Strimmer 2008).

## Acknowledgements

Thanks to Dr. R. Whetten of North Carolina State University, and Dr. J. Schaff of the NCSU Genomic Sciences Lab for their insightful suggestions on ways to improve our methods. Dr. S. Micinski, Dr. J. Lopez, and Dr. J. Westbrook collected the moths used in this project. R. Waples provided one of the custom scripts used in our data pipeline. This project is supported by the Biotechnology Risk Assessment Program competitive grant number 2012-33522-19793 from the USDA - National Institute of Food and Agriculture.

## References

Baeshen, R., Ekechukwu, N., Toure, M., Paton, D., Coulibaly, M., Traoré, S. and Tripet, F. (2014). Differential effects of inbreeding and selection on male reproductive phenotype associated with the colonization and laboratory maintenance of *Anopheles gambiae*. Malar J, 13(1), p.19.

Baxter, S.W., McMillan, O., Chamberlain, N., ffrench-Constant, R.H., Jiggins, C.D. (2009). Prospects for Locating Adaptive Genes in Lepidopteran Genomes. From Molecular Biology and Genetics of the Lepidoptera. CRC Press, Boca Raton, FL, USA. ed. by Goldsmith MR, Marec F. pp. 105–118.

Benjamini, Y., and Hochberg, Y. (1995). Controlling the False Discovery Rate: A practical and powerful approach to multiple testing. Journal of the Royal Statistical Society. Series B (Methodological), 57(1), p.289–300.

Bi, K., Linderoth, T., Vanderpool, D., Good, J., Nielsen, R. and Moritz, C. (2013). Unlocking the vault: next-generation museum population genomics. Molecular Ecology, 22(24), pp.6018–6032.

Blanco, C. (2012). *Heliothis virescens* and Bt cotton in the United States. GM Crops & Food, 3(3), pp.201–212.

Boller, E. (1972). Behavioral aspects of mass-rearing of insects. Entomophaga, 17(1), pp.9–25.

Catchen, J., Amores, A., Hohenlohe, P., Cresko, W., Postlethwait, J. and De Koning, D. (2011). Stacks: Building and Genotyping Loci De Novo From Short-Read Sequences. G3: Genes Genomes Genetics, 1(3), pp.171–182.

Catchen, J., Hohenlohe, P., Bassham, S., Amores, A. and Cresko, W. (2013). Stacks: an analysis tool set for population genomics. Molecular Ecology, 22(11), pp.3124–3140.

Charlesworth, B., Charlesworth, D. (2010). Elements of Evolutionary Genetics. Roberts and Company Publishers, Greenwood, CO, USA. pp. 29.

Charlesworth, D., and Willis, J.H. (2009). The genetics of inbreeding depression. Nature Reviews Genetics, 10, pp. 783–796.

Collard, B., Mace, E., McPhail, M., Wenzl, P., Cakir, M., Fox, G., Poulsen, D., Jordan, D. (2009). How accurate are the marker orders in crop linkage maps generated from large marker datasets? Crop and Pasture Science, 60(4), pp. 362–372.

Collins A.M. Artificial selection of desired characteristics in insects. Advances and Challenges in Insect Rearing, Eds. EG King and NC Leppla. USDA Technical Bulletin. Jan. 1984 pp. 9–19.

Davey, J.W., Cezard, T., Fuentes-Utrilla, P., Eland, C., Gharbi, K., and Blaxter, M.L. (2012). Special features of RAD sequencing data: implications for genotyping. Molecular Ecology, 22, pp. 3151–3164.

Dekker, T., Ibba, I., Siju, K.P., Stensmyr, M.C., Hansson, B.S. (2006). Olfactory shifts parallel superspecialism for toxic fruit in *Drosophila melanogaster* sibling, *D. sechellia*. Current Biology, 16(1), pp. 101–109.

de Valdez, M.F.W., Nimmo, D., Betz, J., Gong, H.F., James, A.A., Alphey, L. and Black, W.C. (2011). Genetic elimination of dengue vector mosquitoes. Proceedings of the National Academy of Sciences, USA, 108(12), pp.4772–4775.

Dobzhansky, T., and Spassky, N. (1954). Environmental modification of heterosis in *Drosophila pseudoobscura*. Proceedings of the National Academy of Sciences, USA, 40(6), pp.407–415.

Elshire, R.J., Glaubitz, J.C., Sun, Q., Poland, J.A., Kawamoto, K., Buckler, E.S., Mitchell, S.E. (2011). A robust, simple genotyping-by-sequencing (GBS) approach for high diversity species. PloS ONE, 6(5), pp. e19379.

Etzel, L.K., and Legner, E.F. (1999). Culture and Colonization, Chapter 7 (pp. 125–176) in Handbook of Biological Control, Academic Press, San Diego, CA, USA. ed. Bellows TS and Fisher TW.

Falconer, D.S., and Mackay TFC (1996) Introduction to Quantitative Genetics. Fourth Edition, Longman Group Ltd. Essex, England. pp. 89.

Fox, C.W., Scheibly, K.L., Smith, B.P., Wallin, W.G. (2007). Inbreeding depression in two seed-feeding beetles *Callosobruchus maculatus* and *Stator limbatus* (Coleoptera: Chrysomelidae). Bulletin of Entomological Research, 97(1), pp.49–54.

Fritz, M.L., Walker, E.D., Miller, J.R., Severson, D.W., Dworkin, I. (2015). Divergent host preferences of above- and below-ground *Culex pipiens* mosquitoes and their hybrid offspring. Medical and Veterinary Entomology, 29(2), p.115–123.

Gahan, L.J., Gould, F., Heckel, D.G. (2001). Identification of a gene associated with Bt resistance in *Heliothis virescens*. Science, 293, pp. 857–860.

Gahan, L.J., Pauchet, Y., Vogel, H., Heckel, D.G. (2010). An ABC transporter mutation is correlated with insect resistance to *Bacillus thuringiensis* Cry1Ac toxin. PloS Genetics, 6(12), pp. e1001248.

Gerloff, C.U., Ottmer, B.K., Schmid-Hempel, B. (2003). Effects of inbreeding on immune response and body size in a social insect, *Bombus terrestris*. Functional Ecology, 17(5), pp.582–589.

Goldman, I. F., Arnold, J., and Carlton, B. C. (1986). Selection for resistance to *Bacillus thuringiensis* subspecies israelensis in field and laboratory populations of the mosquito *Aedes aegypti*. Journal of Invertebrate Pathology, 47(3), pp.317–324.

Gordon, D., Abajian, C., Green, P. (1998). Consed: a graphical tool for sequence finishing. Genome Research, 8, pp. 195–202.

Gould, F., Anderson, A., Reynolds, A., Bumgarner, L., Moar, W. (1995). Selection and genetic analysis of a *Heliothis virescens* (Lepidoptera: Noctuidae) strain with high levels of resistance to *Bacillus thuringiensis* toxins. Journal of Economic Entomology, 88(6), pp.1545–1559.

Groot, A., Gemeno, C., Brownie, C., Gould, F., and Schal, C. (2005). Male and female antennal responses in *Heliothis virescens* and *H. subflexa* to conspecific and heterospecific sex pheromone compounds. Environmental Entomology, 34(2), pp.256–263.

Groot, A.T., Classen, A., Inglis, O., Blanco, C.A., Lopez, J., Teran Vargas, A., Schal, C., Heckel, D.G., and Schofl, G. (2011). Genetic differentiation across North America in the generalist moth *Heliothis virescens* and the specialist *H. subflexa*. Molecular Ecology, 20, pp. 2676–2692.

Hartl, D.L., and Clark, A.G. (2007). Principles of Population Genetics, 4th ed. Sunderland, MA: Sinauer Associates, Inc.

Hoy, M.A. (1990). Genetic improvement of parasites and predators. FFTC-NARC International Seminar on ‘The use of parasitoids and predators to control agricultural pests’, Tukuha Science City, Ibaraki-ken, 305 Japan, October 2-7, 1989. pp.15

Huettel, M.D. (1976). Monitoring the quality of laboratory-reared insects: A biological and behavioral perspective. Environmental Entomology, 5(5), pp.807–814.

Jombart, T. (2008). adegenet: a R package for the multivariate analysis of genetic markers. Bioinformatics, 24, pp. 1403–1405.

Joyner, K. and Gould, F. (1985). Developmental consequences of cannibalism in *Heliothis zea* (Lepidoptera: Noctuidae). Annals of the Entomolgical Society of America, 78, pp. 24–28.

Kajitani, R., Toshimoto, K., Noguchi, H., Toyoda, A., Ogura, Y., Okuno, M., Yabana, M., Harada, M., Nagayasu, E., Maruyama, H., Kohara, Y., Fujiyama, A., Hayashi, T., and Itoh, T. (2014). Efficient d*e novo* assembly of highly heterozygous genomes from whole-genome shotgun short reads. Genome Research, 24, pp. 1384–1395.

Keenan, K., McGinnity, P., Cross, T.F., Crozier, W.W., and Prodöhl, P.A. (2013). diveRsity: An R package for the estimation of population genetics parameters and their associated errors. Methods in Ecology and Evolution, doi: 10.111/2041-210X.12067.

Keller, L.F., and Waller, D.M., (2002). Inbreeding effects in wild populations. Trends in Ecology and Evolution, 17(5), pp.230–241.

Kim, S.H., Cheng, K.M., Ritland, C., Ritland, K., and Silversides, F.G., (2007). Inbreeding in the Japanese quail estimated by pedigree and microsatellite analyses. Journal of Heredity, 98(14), pp. 378–381.

Kosambi, D.D. (1944). The estimation of map distance from recombination values. Annals of Eugenics, 12, pp. 172–175.

Li, R., Li, Y., Fang, X., Yang, H., Wang, J., Kristiansen, K., and Wang, J. (2009). SNP detection for massively parallel whole-genome resequencing. Genome Research, 19, pp. 1124–1132.

Mackauer, M. (1976). Genetic problems in the production of biological control agents. Annual Review of Entomology, 21, pp. 369–385.

Mackay, T.F.C., Richards, S., Stone, E.A., Barbadilla, A., Ayroles, J.F., Zhu, D., et al., (2012) The *Drosophila melanogaster* genetic reference panel. Nature, 482, pp. 173–178.

Magoc, T., and Salzberg, S. (2011). FLASH: Fast length adjustment of short reads to improve genome assemblies. Bioinformatics, 27(21), pp.2957–2963.

Margarido, G. R. A., Souza, A. P. and Garcia, A. A. F. (2007). OneMap: software for genetic mapping in outcrossing species. Hereditas, 144, pp. 78–79.

Mita, K., Kasahara, M., Sasaki, S., Nagayasu, Y., Yamada, T., Kanamori, H., Namiki, N., Kitagawa, M., Yamashita, H., Yasukochi, Y., Kadono-Okuda, K., Yamamoto, K., Ajimura, M., Ravikumar, G., Shimomura, M., Nagamura, Y., Shin-i, T., Abe, H., Shimada, T., Morishita, S., and Sasaki, T. (2004) The Genome Sequence of Silkworm, *Bombyx mori*. DNA Research, 11(1), pp. 27–35

Mollinari, M., Margarido, G.R., Vencovsky, R., Garcia, A.A. (2009). Evaluation of algorithms used to order markers on genetic maps. Heredity, 103(6), p.494–502.

Mukhopadhyay, J., Rangel, E.F., Ghosh, K., Munstermann, L.E. (1997). Patterns of genetic variability in colonized strains of *Lutzomyia longipalpis* (Diptera: Psychodidae) and its consequences. American Journal of Tropical Medicine and Hygiene, 57(2), pp.216–221.

Munstermann, L.E. (1994). Unexpected genetic consequences of colonization and inbreeding: allozyme tracking in Culicidae (Diptera). Annals of the Entomological Society of America, 87(2), pp.157–164.

Nei M (1987) Molecular evolutionary genetics. New York: Columbia University Press.

Nickerson, D.A., Tobe, V.O., and Taylor, S.L. (1997). PolyPhred: automating the detection and genotyping of single nucleotide substitutions using fluorescence-based resequencing. Nucleic Acids Research, 25(14), pp.2745–2751.

Nielsen, R., Paul, J.S., Albrechtsen, A., Song, Y.S. (2011). Genotype and SNP calling from next-generation sequencing data. Nature Reviews Genetics, 12, pp. 443–451.

Norris, D.E., Shurtleff, A.C., Toure, Y.T., Lanzaro, G.C. (2001). Microsatellite DNA polymorphism and heterozygosity among field and laboratory populations of *Anopheles gambiae* s.s. (Diptera: Culicidae). Journal of Medical Entomology, 38(2), pp.336–340.

Oppenheim, S.J., Gould, F., Hopper, K.R. (2012). The genetic architecture of a complex ecological trait: host plant use in the specialist moth, *Heliothis subflexa*. Evolution, 66(11), pp. 3336–3351.

Paradis, E. (2010). pegas: an R package for population genetics with an integrated-modular approach. Bioinformatics, 26, 419–420.

Paterson, A.H. (1996). Making genetic maps. A.H. Paterson (Ed.), Genome Mapping In Plants, R. G. Landes Company, San Diego, California, pp. 23–29

Peterson, B.K., Weber, J.N., Kay, E.H., Fisher, H.S., Hoekestra, H.E. (2012). Double digest RADseq: An inexpensive method of *de novo* SNP discovery and genotyping in model and non-model species. PloS ONE, 7(5), e37135.

Poland, J.A., Brown, P.H., Sorrells, M.E., Jannink, J.L. (2012). Development of high-density genetic maps for barley and wheat using a novel two-enzyme genotyping-by-sequencing approach. PLoS ONE, 7(2), pp. e32253.

Pootakham, W., Jomchai, N., Ruang-areerate, P., Shearman, J. R., Sonthirod, C., Sangsrakru, D. Tragoonrun, S., Tangphatsornruang, S., (2015). Genome-wide SNP discovery and identification of QTL associated with agronomic traits in oil palm using genotyping-by-sequencing (GBS). Genomics, 105(5), pp.288–295.

Pradeep, A. R., Chatterjee, S. N., & Nair, C. V. (2005). Genetic differentiation induced by selection in an inbred population of the silkworm *Bombyx mori*, revealed by RAPD and ISSR marker systems. Journal of Applied Genetics, 46(3), pp. 291.

R core team (2014). R: A language and environment for statistical computing. R Foundation for Statistical Computing, Vienna, Austria. URL http://www.R-project.org/.

Raulston, J.R. (1975). Tobacco budworm: observations on the laboratory adaptation of a wild strain. Annals of the Entomological Society of America, 68, pp. 139–142.

Roush, R.T. (1986). Inbreeding depression and laboratory adaptation in *Heliothis virescens* (Lepidoptera: Noctuidae). Annals of the Entomological Society of America, 79, pp. 583–587.

Rumball, W., Franklin, I., Frankham, R., Sheldon, B. (1994). Decline in heterozygosity under full-sib and double first-cousin inbreeding in *Drosophila melanogaster*. Genetics, 136, pp. 1039–1049.

Shaw, K.L. (2000). Interspecific genetics of mate recognition: inheritance of female acoustic preference in Hawaiian crickets. Evolution, 54(4), pp.1303–1312.

Sheck, A. L., and Gould, F. (1995). Genetic analysis of differences in oviposition preferences of *Heliothis virescens* and *H. subflexa* (Lepidoptera: Noctuidae). Environmental Entomology, 24(2), p.341–347.

Sheck, A. L., and Gould, F. (1996). The genetic basis of differences in growth and behavior of specialist and generalist herbivore species: selection on hybrids of *Heliothis virescens* and *Heliothis subflexa* (Lepidoptera). Evolution, 50(2), pp.831–841.

Sheck, A. L., Groot, A. T., Ward, C. M., Gemeno, C., Wang, J., Brownie, C., Schal, C., and Gould, F. (2006). Genetics of sex pheromone blend differences between *Heliothis virescens* and *Heliothis subflexa*: a chromosome mapping approach. Journal of Evolutionary Biology, 19(2), pp. 600–617.

Singh, R., Tan, S.G., Panandam, J.M., Rahman, R.A., Ooi, L.C.L., Low, E.L., Sharma, M., Jansen, J., and Cheah, S. (2009). Mapping quantitative trait loci (QTLs) for fatty acid composition in an interspecific cross of oil palm. BMC Plant Biology, 9, 114.

Sokal, R.R., and Rohlf, F.J. (1995). Biometry: The Principles and Practices of Statistics in Biological Research. 3rd. ed. New York: W.H. Freeman and Company.

Sokolowski, M.B. (1980). Foraging strategies of *Drosophila melanogaster*: A chromosomal analysis. Behavioral Genetics, 10(3), p.291–302.

Strimmer, K. (2008). fdrtool: a versatile R package for estimating local and tail area-based false discovery rates. Bioinformatics Applications, 24(12), pp.1461–1462.

Tajima, F. (1989) Statistical method for testing the neutral mutation hypothesis by DNA polymorphism. Genetics, 123, pp. 585–595.

Taylor, M.F.J., Heckel, D.G., Brown, T.M., Kreitman, M.E., Black, G. (1993). Linkage of pyrethroid resistance to a sodium channel locus in the tobacco budworm. Insect Biochemistry and Molecular Biology, 23(7), pp.763–775.

Tomaru, M., Doi, M., Higuchi, H., Oguma, Y. (2000). Courtship song recognition in the *Drosophila melanogaster* complex: heterospecific songs make females receptive in *D. melanogaster*, but not in *D. sechellia*. Evolution, 54(4), pp.1286–1294.

Turissini, D.A., Gamez, S., White, B.J. (2014). Genome-wide patterns of polymorphism in an inbred line of the African malaria mosquito, *Anopheles gambiae*. Genome Biology and Evolution, 6(11), p.3094–3104.

Van Os, H., Stam, P., Visser, R.G.F., Van Eck, H.J. (2005). RECORD: a novel method for ordering loci on a genetic linkage map. Theoretical and Applied Genetics, 112, pp. 30–40.

Ward, M.D. (2009). Genetics of Sex Pheromones: Mapping Desaturase Genes in *Heliothis* Species (Unpublished Master’s Thesis). North Carolina State University, Raleigh, NC.

Watterson, G.A. (1975). On the number of segregating sites in genetical models without recombination. Theoretical Population Biology, 7, pp. 256–276.

Weir, B.S., and Cockerham C.C. (1984). Estimating F-statistics for the analysis of population structure. Evolution, 38(6), p.1358–1370.

Xu, P., Xu, S., Wu, X., Tao, Y., Wang, B., Wang, S., Qin, D., Lu, Z., Li, G. (2014). Population genomic analyses from low-coverage RAD-Seq data: a case study on the non-model cucurbit bottle gourd. The Plant Journal, 77, pp. 430–442.

You, M., Yue, Z., He, W., Yang, X., Yang, G., Xie, M., Zhan, D., Baxter, S.W., et al. (2013). A heterozygous moth genome provides insights into herbivory and detoxification. Nature Genetics, 45, pp. 220–225.

